# Neurotoxins subvert the allosteric activation mechanism of SARM1 to induce neuronal loss

**DOI:** 10.1101/2021.07.16.452689

**Authors:** Tong Wu, Jian Zhu, Amy Strickland, Kwang Woo Ko, Yo Sasaki, Caitlin Dingwall, Yurie Yamada, Matthew D Figley, Xianrong Mao, Alicia Neiner, Joseph Bloom, Aaron DiAntonio, Jeffrey Milbrandt

**Affiliations:** Department of Genetics, Washington University Medical School, St. Louis, MO 63110, USA; Department of Developmental Biology, Washington University Medical School, St. Louis, MO 63110, USA; Needleman Center for Neurometabolism and Axonal Therapeutics, Washington University School of Medicine in Saint Louis, St. Louis, MO 63114, USA; Department of Biomedical Engineering, Washington University in St. Louis, St. Louis, MO 63130, USA

## Abstract

SARM1 is an inducible TIR-domain NAD^+^ hydrolase that mediates pathological axon degeneration. SARM1 is activated by an increased ratio of NMN to NAD^+^, which competes for binding to an allosteric activating site. When NMN binds, the TIR domain is released from autoinhibition, activating its NAD^+^ hydrolase activity. The discovery of this allosteric activating site led us to hypothesize that other NAD^+^-related metabolites might also activate SARM1. Here we show that the nicotinamide analogue 3-acetylpyridine (3-AP), first identified as a neurotoxin in the 1940s, is converted to 3-APMN which activates SARM1 and induces SARM1-dependent NAD^+^ depletion, axon degeneration and neuronal death. Systemic treatment with 3-AP causes rapid SARM1-dependent death, while local application to peripheral nerve induces SARM1-dependent axon degeneration. We also identify a related pyridine derivative, 2-aminopyridine, as another SARM1-dependent neurotoxin. These findings identify SARM1 as a candidate mediator of environmental neurotoxicity, and furthermore, suggest that SARM1 agonists could be developed into selective agents for neurolytic therapy.

## INTRODUCTION

Axon degeneration is an essential feature of many neurodegenerative conditions. SARM1 is the central executioner of pathological axon degeneration, and a promising therapeutic target for a wide range of neurological disorders (DiAntonio, 2019). SARM1 is comprised of an autoinhibitory N-terminal ARM domain, tandem SAM domains that mediate octamerization, and a C-terminal TIR domain. This TIR domain is the original exemplar of a large family of TIR domain NAD^+^ hydrolases, conserved throughout all domains of life from bacteria and archaea to plants and animals (Essuman et al., 2017, 2018; Horsefield et al., 2019; Wan et al., 2019). In healthy neurons, SARM1 is maintained in an autoinhibited state via multiple intra- and intermolecular interactions (Shen et al., 2021), including binding of the N-terminal ARM domain to the C-terminal TIR-domain (Summers et al., 2016). SARM1 autoinhibition is regulated by an allosteric binding site within the autoinhibitory ARM domain that can bind either nicotinamide adenine dinucleotide (NAD^+^) (Jiang et al., 2020; Sporny et al., 2020) or its precursor, nicotinamide mononucleotide (NMN) (Figley et al., 2021). Axon injury leads to loss of the NAD^+^ biosynthetic enzyme NMNAT2 (Gilley et al., 2010), resulting in an increased NMN/NAD^+^ ratio that favors NMN binding to the allosteric site (Figley et al., 2021). The switch from NAD^+^ to NMN binding induces compaction of the autoinhibitory ARM domain and permits the formation of TIR-TIR interactions that activate the enzyme (Figley et al., 2021). The identification of this allosteric binding pocket led us to hypothesize that metabolites with structural similarities to NMN and NAD^+^ might also regulate SARM1 activation.

To explore whether additional NAD^+^-related metabolites activate SARM1, we searched the literature for neurotoxins with proposed mechanisms of action related to NAD^+^ metabolism. NAD^+^ biosynthesis in mammals largely occurs via the formation of NMN from nicotinamide (Nam a.k.a. vitamin B3) by nicotinamide phosphoribosyl transferase (NAMPT), followed by NAD^+^ synthesis from NMN, catalyzed by a nicotinamide mononucleotide adenyltransferase (NMNAT) enzyme (Revollo et al., 2004; Wang et al., 2006) (Fig. 1a). Studies of anti-metabolites starting in the 1940s demonstrated that 3-acetylpyridine (3-AP), a Nam analogue, is a neurotoxin that causes degeneration in both the central and peripheral nervous systems (Beher et al., 1952; Desclin et al., 1974; Hicks, 1955; Lopiano et al., 1986; Schulz et al., 1994; Wecker et al., 2017; Woolley et al., 1943). 3-AP is structurally identical to Nam except that a methyl group replaces Nam’s amino group. Interestingly, 3-AP neurotoxicity is prevented by co-administration of Nam. This led to the hypothesis that 3-AP toxicity is due to the substitution of 3-AP for Nam in the NAD^+^ biosynthetic pathway, generating 3-acetylpyridine adenine dinucleotide (3-APAD) (Fig. 1a), a potential NAD^+^ mimetic that subsequently disrupts NAD^+^-dependent metabolic processes. While 3-AP is converted to 3-APAD, it is not clear why neurons would be particularly sensitive to this toxin. Severe vitamin B3 deficiency, i.e. pellagra, decreases NAD^+^ levels and results in abnormalities in many tissues throughout the body. We therefore hypothesized that 3-AP introduction into the NAD^+^ biosynthesis pathway causes neurodegeneration not via defects in NAD^+^ biosynthesis or 3-APAD interference in NAD^+^-regulated processes, but rather through inappropriate SARM1 activation.

**Figure 1.**
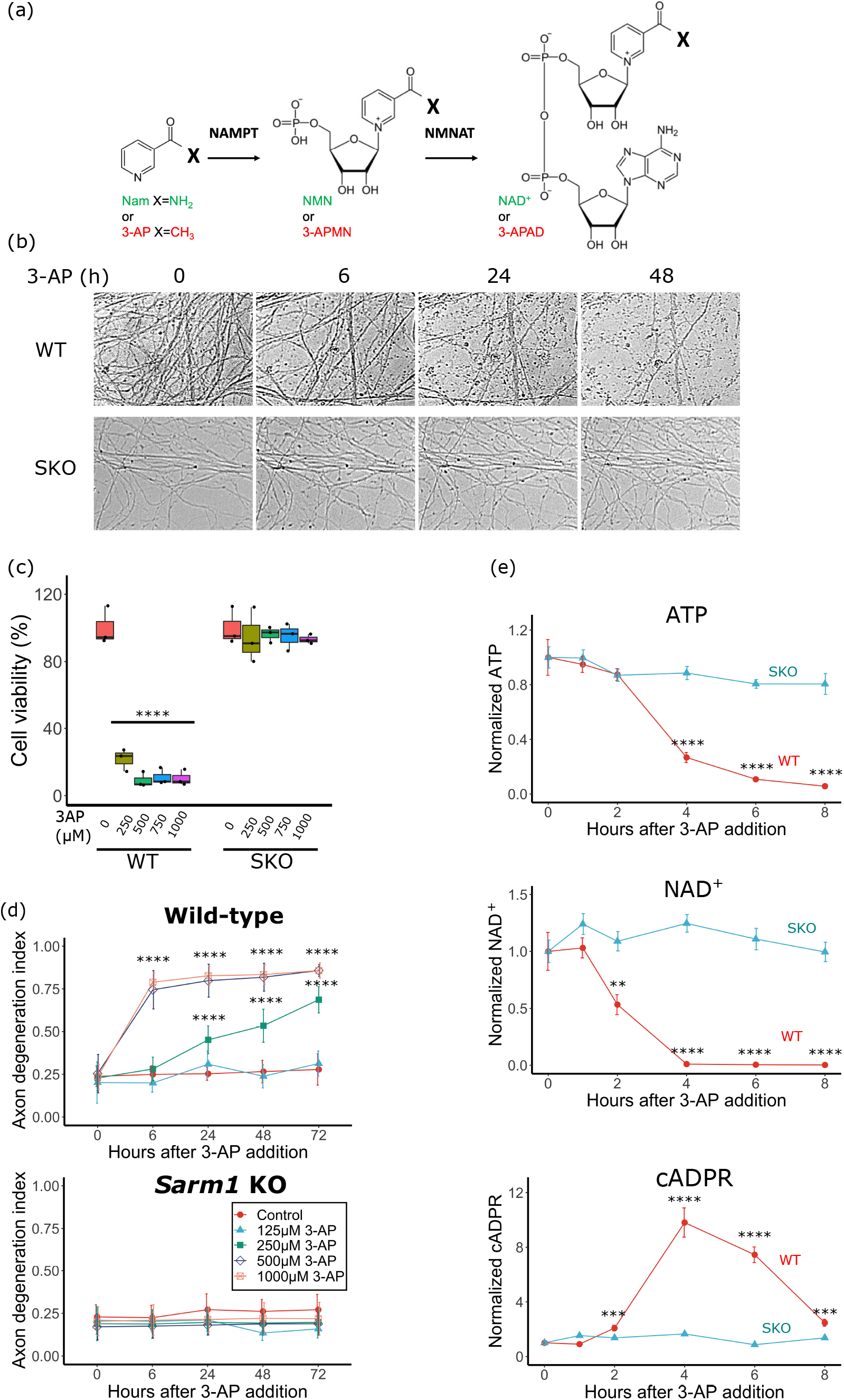
3-AP causes SARM1-dependent neurotoxicity. (a) Diagram of nicotinamide (Nam) and 3-AP metabolism by NAMPT and NMNAT. X represents amino (Nam) or methyl (3-AP) group. (b) WT and *Sarm1* KO DRGs were treated with 3-AP (300 μM) at DIV7. Representative images of axons from WT and *Sarm1* KO cultured DRG neurons at 0, 6, 24, 48 hr after 3-AP treatment show axon degeneration. (c) WT and *Sarm1* KO embryonic DRGs were treated with 3-AP for 12 hr and MTT assays were performed to quantify cell viability at DIV7. The first and third quartile, and the median are shown in boxplot with ± 1.5 time interquartile. Statistical significance was determined by one-way ANOVA test, comparing to cell viability of each genotype treated with 0 μM 3-AP. (d) WT (top) and *Sarm1* KO (bottom) DRGs were treated with 3-AP (300 μM) at DIV7. Axon degeneration index was measured at indicated times after 3-AP treatment. Data with error bars correspond to Mean±SD. Statistical significance was determined by one-way ANOVA tests, comparing to control degeneration index at time 0 hr. (e) WT or *Sarm1* KO DRG neurons were treated with 3-AP at DIV7. Metabolites were quantified by LC-MS/MS at indicated times after treatment. Data with error bars correspond to Mean±SD. Statistical significance was determined by one-way ANOVA tests, comparing to metabolite of each genotype at time 0 hr. *p < 0.05; **p < 0.01; ***p < 0.001; ****p < 0.0001.

To test this hypothesis, we investigated the impact of 3-AP on neurons both *in vitro* and *in vivo*. We find that incubation of cultured mouse dorsal root ganglion (DRG) neurons with 3-AP leads to SARM1 activation, and SARM1-dependent NAD^+^ depletion, axon degeneration, and cell death. This neurotoxicity requires both the conversion of 3-AP to 3-acetylpyridine mononucleotide (3-APMN), and a properly-formed SARM1 allosteric NMN-binding site. We also investigated the impact of 3-AP *in vivo* and found that systemic exposure to 3-AP induces rapid death in wild type (WT) mice, but causes no obvious harm to *Sarm1* knockout (KO) mice. Local application of 3-AP to peripheral nerves leads to SARM1-dependent axon degeneration. Finally, we assayed additional nicotinamide analogs and found that 2-aminopyridine also triggers SARM1-dependent neurodegeneration. These findings identify SARM1 as a target of multiple neurotoxins and a candidate mediator of environmental neurotoxicity in neurodegenerative disease. In addition, the identification of exogenous chemical SARM1 agonists may enable the development of new methods for neurolytic therapy.

## RESULTS

### 3-AP induces SARM1-dependent axon degeneration and neuron death

3-acetyl-pyridine (3-AP) is an analog of nicotinamide (Nam) in which the amino group is replaced by a methyl group (Fig. 1a). 3-AP was previously studied as an anti-metabolite in NAD^+^ biosynthesis, and assumed to produce its effects by disrupting the conversion of Nam to NMN and ultimately NAD^+^. Early studies showed that administration of 3-AP to rabbits, dogs, mice and rats resulted in severe neurotoxicity and death that could be blocked by administration of Nam (Jolicoeur et al., 1982; Weller et al., 1992).

The recent demonstration that NMN binds to an allosteric site in SARM1 to activate its NAD^+^ hydrolase activity and promote axon degeneration led us to examine the role of SARM1 in 3-AP toxicity. We cultured WT mouse embryonic dorsal root ganglion (DRG) neurons and treated them with varying doses of 3-AP. We measured axonal integrity at multiple time points after 3-AP addition and find that axons remain intact at low doses, but begin to degenerate slowly when exposed to 250 μM 3-AP. At doses above 250 μM, axons rapidly degenerate at a rate similar to that observed after transection (Fig. 1b, d). We also measured neuronal death using the MTT assay and find that 3-AP treatment causes neuronal death at similar doses (Fig. 1c).

SARM1 is required for pathological axon degeneration downstream of multiple insults (Figley & DiAntonio, 2020). We tested whether 3-AP-mediated axon degeneration and neuronal death also require SARM1 by treating *Sarm1* KO DRG neurons with escalating doses of 3-AP. Surprisingly, we find that 3-AP has no effect on neurons lacking SARM1 (Fig. 1c, d), even at extremely high doses (e.g., 1 mM). These experiments demonstrate that 3-AP mediated neurotoxicity requires SARM1.

When SARM1 is activated by axonal injury, its potent NAD^+^ hydrolase activity depletes the axon of NAD^+^ and produces cADPR, a sensitive biomarker of SARM1 activity (Essuman et al., 2017; Sasaki et al., 2020). These metabolic changes incite energetic catastrophe and subsequent axon fragmentation (Gerdts et al., 2015). To determine whether 3-AP activates SARM1 NAD^+^ hydrolase activity, we measured relevant metabolites in neurons following addition of 3-AP (300 μM). We find substantial NAD^+^ and ATP depletion and a ~10-fold increase in cADPR at 4 hr post 3-AP addition (Fig. 1e). These metabolite changes are similar to those observed after axotomy and indicate that SARM1 is activated by 3-AP. To confirm the SARM1-dependence of 3-AP neurotoxicity, we performed similar metabolite measurements in 3-AP-treated *Sarm1* KO neurons and find no significant changes in the levels of NAD^+^, ATP, or cADPR (Fig. 1e). Taken together, these results demonstrate that 3-AP stimulates metabolite changes consistent with SARM1 activation, and that 3-AP neurotoxicity is dependent on SARM1.

### Neurotoxicity caused by 3-AP requires its conversion to 3-APMN

While the metabolic changes induced by 3-AP are almost identical to those caused by axotomy, there is one telling difference. Following axotomy, NMN levels rise because axonal NMNAT2 is lost (Di Stefano et al., 2015). In contrast, 3-AP intoxication leads to a greater than 50% decline in NMN within four hours (p<0.001, n=3). As NMN is synthesized from Nam and PRPP by the enzyme NAMPT, this finding suggests that 3-AP competes with Nam for NAMPT, and thereby inhibits the production of NMN. Early studies of 3-AP neurotoxicity in mice and other species provide additional evidence for competition between 3-AP and Nam. These studies showed that co-administration of Nam prevented 3-AP-induced neurotoxicity and death (Jolicoeur et al., 1982; Weller et al., 1992). To assess whether Nam influences 3-AP-mediated axon degeneration, we pretreated DRG neurons with increasing amounts of Nam before addition of 3-AP. We find that Nam pretreatment reduces 3-AP-induced axon degeneration in a dose-dependent manner (Fig. 2a), suggesting that Nam blocks the metabolism of 3-AP into a compound toxic to axons. As an independent test of this hypothesis, we treated neurons with 3-AP and FK866, an inhibitor of NAMPT. NAMPT inhibition blocks the neurotoxic actions of 3-AP (Fig. 2b), supporting the idea that 3-AP is metabolized to a toxic derivative via a pathway involving NAMPT.

**Figure 2.**
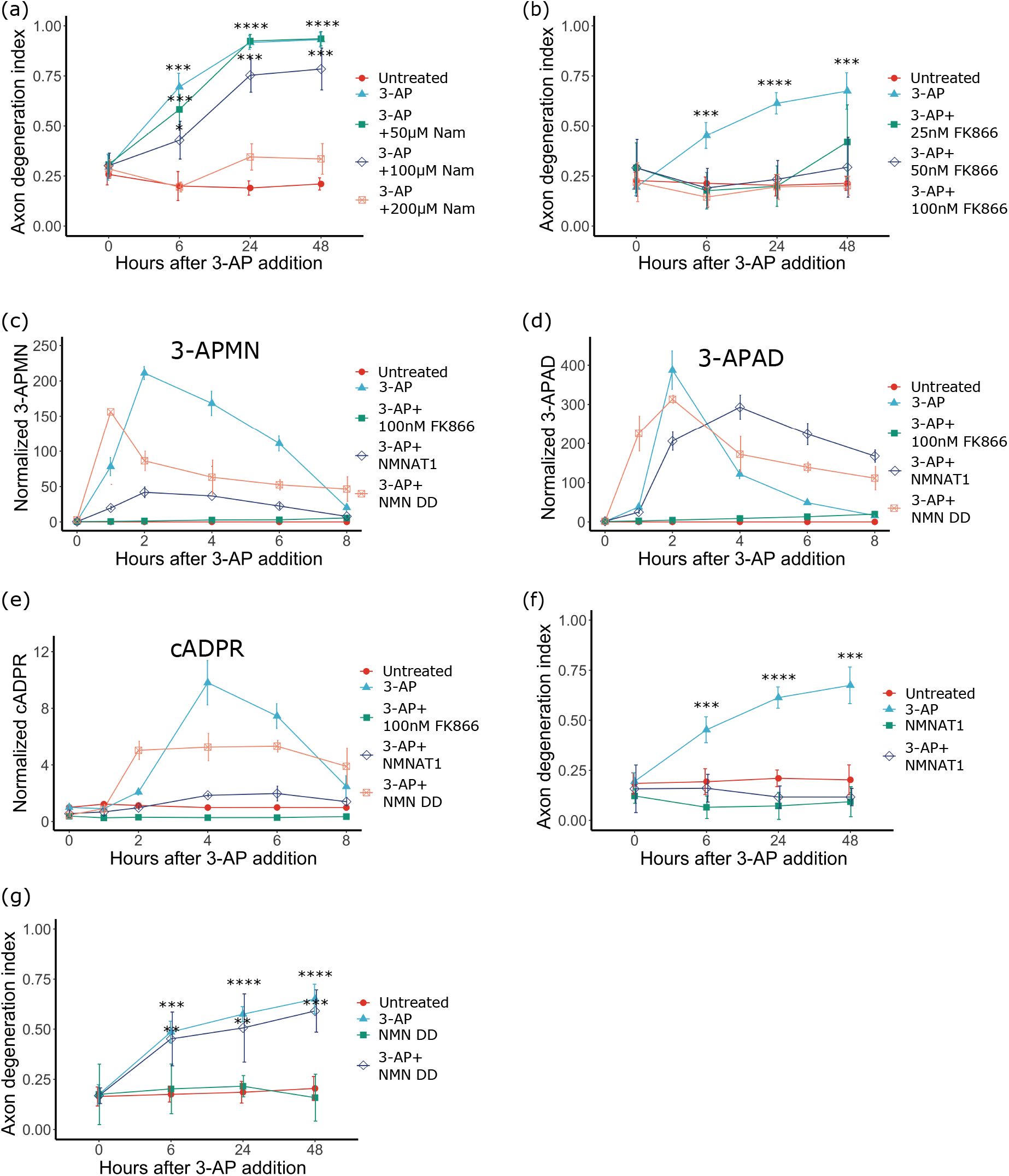
The conversion of 3-AP to 3-APMN is required for neurotoxicity. (a) Nam inhibits 3-AP-induced axon degeneration. WT DRG neurons were pretreated overnight with Nam at 0, 50, 100 or 200 μM on DIV7, then treated with 3-AP (500 μM). Axon degeneration was quantified at indicated times following treatment. Statistical significance was determined by one-way ANOVA tests (compared to 0 hr). (b) NAMPT inhibitor FK866 protects neurons against 3-AP toxicity. WT DRG neurons were pretreated overnight with FK866 at 0, 25, 50 or 100 nM, then treated with 3-AP (300 μM). Axon degeneration was quantified after 3-AP addition. Statistical significance was determined by one-way ANOVA test (compared to 0 hr). (c-e) Metabolites in WT neurons treated as indicated were measured using LC-MS/MS. Statistical significance was determined by one-way ANOVA test (compared to 0 hr). (f) NMNAT1 prevents 3-AP-induced axon degeneration. WT DRG neurons were infected with NMNAT1 lentivirus at DIV3 and treated with 3-AP (300 μM) at DIV7. Statistical significance was determined by one-way ANOVA test (compared to 0 hr). (g) NMN deamidase fails to prevent 3-AP-induced axon degeneration. WT DRG neurons were infected with NMN deamidase (NMN DD) lentivirus at DIV3 and treated with 3-AP (300 μM) at DIV7. Statistical significance was determined by oneway ANOVA test (compared to 0 hr). *p < 0.05; **p < 0.01; ***p < 0.001; ****p < 0.0001. Data with error bars correspond to Mean±SD.

To identify the 3-AP derivative responsible for SARM1 activation and axon degeneration, we first developed mass spectrometry methods to detect 3-acetylpyridine mononucleotide (3-APMN) and 3-acetylpyridine adenine dinucleotide (3-APAD) in neurons (Supplemental Fig. 1a, b). We isolated metabolites from DRG neurons at various timepoints following 3-AP treatment and analyzed them by LC-MS/MS. Both 3-APMN and 3-APAD levels peak within 2 hr of 3-AP addition, with 3-APMN detectable slightly earlier than 3-APAD consistent with a precursor-product relationship (Fig. 2c, d). Notably, they both appear several hours before the increase of the SARM1-derived product cADPR (Fig. 2e).

To demonstrate directly the involvement of NAMPT/NMNAT activity in the production of these metabolites, we pre-treated neurons with FK866 prior to 3-AP addition. In FK866-treated neurons, 3-APMN and 3-APAD remain at baseline, and cADPR levels do not increase (Fig. 2c-e). Together with results demonstrating FK866 blocks axon degeneration in 3-AP treated neurons, these data strongly support the hypothesis that 3-AP is metabolized by NAMPT alone or in combination with other NAD^+^ biosynthetic enzymes (e.g., NMNAT) to produce a SARM1 agonist.

SARM1-mediated axon degeneration caused by mechanical or chemical insults can be blocked by overexpression of NMNAT1 (Araki et al, 2004) or the bacterial enzyme NMN deamidase (Di Stefano et al., 2015). The axon protection provided by these enzymes presumably reflects their ability to reduce axonal NMN levels via conversion of NMN to NAD^+^ or NaMN, respectively. When NMN levels are low, it is less likely to bind the SARM1 allosteric site and induce the conformational change that activates SARM1 hydrolase activity and subsequent axon degeneration. To investigate which 3-AP-derived metabolite activates SARM1, we tested whether these NMN-consuming enzymes could also ameliorate 3-AP neurotoxicity. We find that NMNAT1 prevents 3-AP-induced increases in cADPR and axon fragmentation (Fig. 2e, f). In contrast, axons of neurons expressing NMN deamidase remain susceptible to 3-AP and high levels of cADPR are produced (Fig. 2e, g). Importantly, 3-APMN and 3-APAD are still generated in the presence of NMN deamidase (Fig. 2c, d). This is consistent with its deamidase activity, as 3-AP derivatives lack the amino group that is the target of this enzyme in Nam. In contrast, NMNAT1 overexpression strongly blunts the rise of both 3-APMN and cADPR while maintaining robust production of 3-APAD (Fig. 2c-e), indicating that SARM1 is not activated when 3-APMN levels are low. These results support the hypothesis that 3-APMN is the 3-AP derivative that directly activates SARM1.

### 3-AP can be converted to 3-APMN and 3-APAD by NAMPT and NMNAT

The inhibition of NAMPT and overexpression of NMNAT1 both prevent 3-AP mediated SARM1 activation and neuronal toxicity. These results along with the structural similarities between Nam and 3-AP and their respective metabolites, NMN and 3-APMN, led us to hypothesize that NAMPT converts 3-AP to 3-APMN, which then acts as an NMN mimetic to directly activate SARM1-mediated axon degeneration. We first tested whether NAMPT can convert 3-AP to 3-APMN using an *in vitro* biochemical assay. NAMPT enzyme was produced in *E. coli* and the purified enzyme was incubated with 3-AP and its cofactors phosphoribosyl pyrophosphate (PRPP) and ATP. We used LC-MS/MS to detect a novel compound generated in this reaction, which was determined to be 3-APMN (Supplemental Fig. 1a). We next examined whether this 3-APMN could be converted to 3-APAD by NMNAT1. We produced and purified NMNAT1 and performed reactions in which NAMPT and NMNAT1 together were combined with 3-AP, PRPP and ATP. We detected a compound that was identified as 3-APAD using a 3-APAD standard and LC-MS/MS (Fig. 3a; Supplemental Fig. 1b). The production and identification of these two 3-AP derived products confirms that 3-AP can be metabolized by the NAMPT/NMNAT pathway in a parallel fashion to Nam.

**Figure 3.**
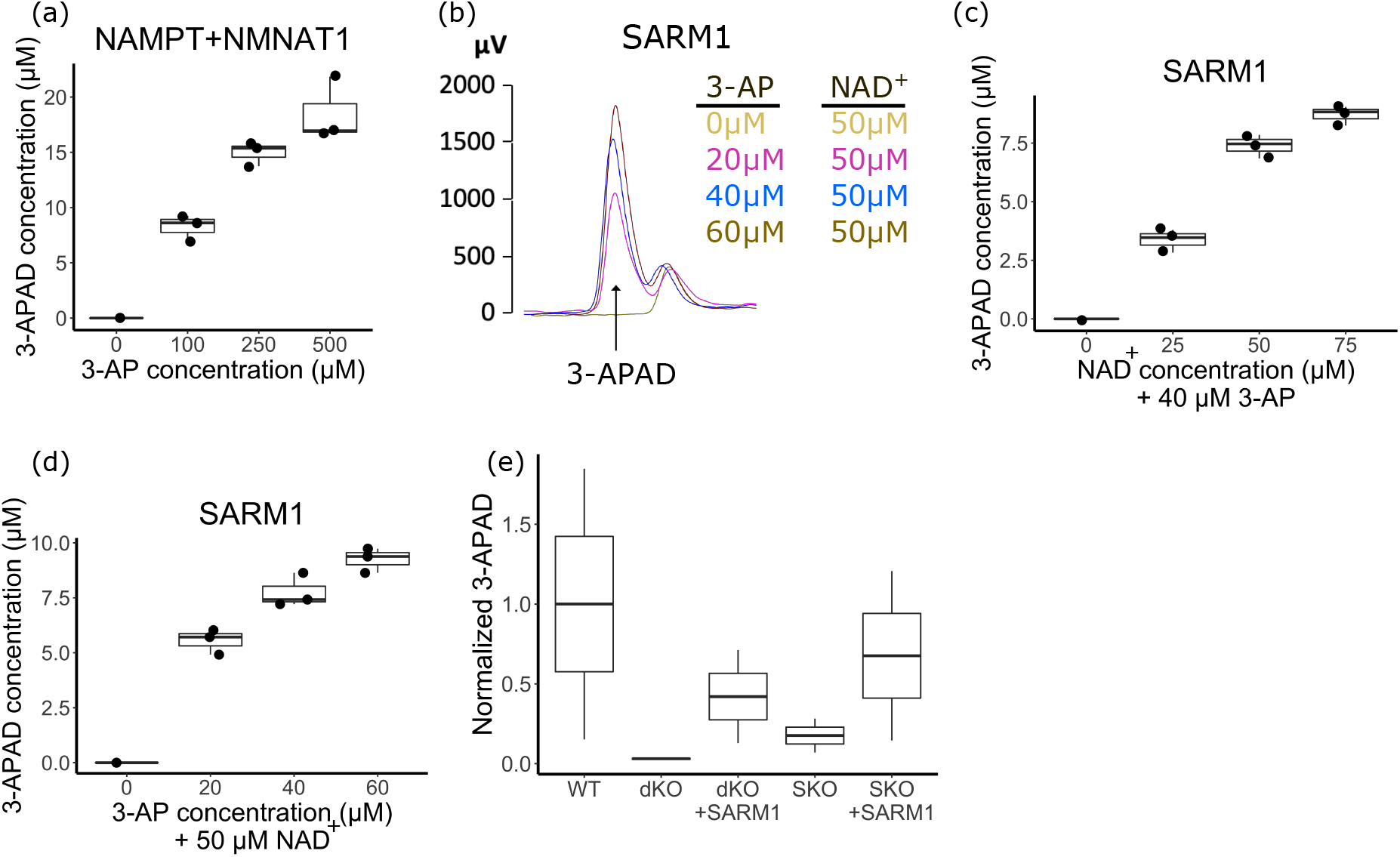
3-APAD can be generated by either NAMPT-NMNATpathway or SARM1-mediated base exchange reaction. (a) 3-APAD is synthesized in 3-AP dose-dependent manner using in vitro reactions containing purified NAMPT and NMNAT1. (b) Representative HPLC trace demonstrating the 3-AP dose-dependent production of 3-APAD in vitro via SARM1-mediated base-exchange reaction. (c) In vitro 3-APAD production by SARM1 is dependent on NAD^+^ concentration at a fixed 3-AP concentration (40 μM). (d) In vitro 3-APAD production by SARM1 is dependent on 3-AP concentration at fixed NAD^+^ concentration (50 μM). (e) 3-APAD in neurons is produced by both SARM1 and the NAMPT/NMNAT pathway. WT, *Sarm1* KO (SKO) and *Nmnat2/ Sarm1* double knockout (dKO) neurons were treated with 3-AP (300 μM) and metabolites were analyzed after 4 hr of treatment. Infection of dKO or SKO neurons with a SARM1 lentivirus increases the amount of 3-APAD detected. The first and third quartile, the median are shown in boxplot with ± 1.5 time interquartile.

### SARM1 also synthesizes 3-APAD from 3-AP and NAD^+^ via base exchange

SARM1 possesses base exchange activity in addition to its hydrolase activity (Zhao et al., 2019), in which the ADPR moiety of NAD^+^ is condensed with an acceptor molecule. This reaction presents a pathway for the direct production of 3-APAD by SARM1 using 3-AP as the acceptor. To test this idea, we performed biochemical assays using SARM1 purified from HEK 293T cells. SARM1 was incubated with NAD^+^ and 3-AP and the reaction products were analyzed using HPLC and LC-MS/MS. We detected the formation of 3-APAD by SARM1 (Fig. 3b) that required both NAD^+^ and 3-AP with the amount of product dependent on the concentrations of both precursors. (Fig. 3c, d). The formation of 3-APAD from 3-AP is consistent with the SARM1 catalyzed transfer of the NAD^+^ ADPR moiety to 3-AP via a base exchange reaction (Zhao et al., 2019). This represents a second route for production of 3-APAD from 3-AP that is in addition to the NAMPT/ NMNAT1 pathway (Fig. 3a).

To determine whether this SARM1-mediated base exchange reaction occurs in neurons, we sought to analyze the generation of 3-APAD in neurons lacking the NAMPT/NMNAT pathway. Because *Nmnat2* knockout mice are not viable (Gilley & Coleman, 2010), we used DRG neurons from *Sarm1/Nmnat2* double-knockout (dKO) mice for these experiments. We find that *Sarm1/Nmnat2* dKO neurons treated with 3-AP do not produce 3-APAD. This demonstrates that 3-APAD production in cultured DRG neurons requires NMNAT2, but does not require its paralogs NMNAT1 or NMNAT3 (Fig. 3e). To further examine the import of the SARM1-mediated base exchange reaction, we infected *Sarm1*/*Nmnat2* dKO neurons with lentivirus expressing SARM1 at DIV3 and examined metabolites at DIV7. We find that re-introducing SARM1 to these neurons restores 3-APAD production (Fig. 3e), demonstrating that SARM1 carries out the base exchange reaction in neurons. Indeed, *Sarm1* KO neurons treated with 3-AP produce much less 3-APAD than do wild type neurons (Fig. 3e), indicating that SARM1 is a major contributor to 3-APAD synthesis. The residual 3-APAD produced in *Sarm1* KO neurons treated with 3-AP demonstrates that the NAMPT/NMNAT pathway also actively produces 3-APAD from 3-AP. Together these results show that both SARM1-catalyzed base exchange activity and NAMPT/NMNAT2 contribute to 3-APAD synthesis in neurons.

### 3-APMN activates purified SARM1 *in vitro*

The results from manipulating the NAMPT/NMNAT pathway and SARM1 in neurons support the hypothesis that 3-APMN can activate SARM1. To test whether SARM1 is directly activated by 3-APMN, we purified SARM1 on Strep-tag beads and incubated it with purified 3-APMN (Supplemental Fig. 2) in the presence of the pyridine conjugate PC6, a fluorescent probe of SARM1 hydrolase activity (Li et al., 2021; Zhao et al., 2019). We measured the reaction in real time by following the fluorescence output signal and found that SARM1 hydrolase activity increases with 3-APMN concentration (Fig. 4a). We determined the activation constant of the reaction (K_A_) to be ~44 μM, which is slightly lower than the reported dissociation constant (K_D_) for NAD^+^ binding to the SARM1 ARM domain (Figley et al., 2021). These results indicate that 3-APMN interacts directly with SARM1 to activate its NAD^+^ hydrolase activity.

**Figure 4.**
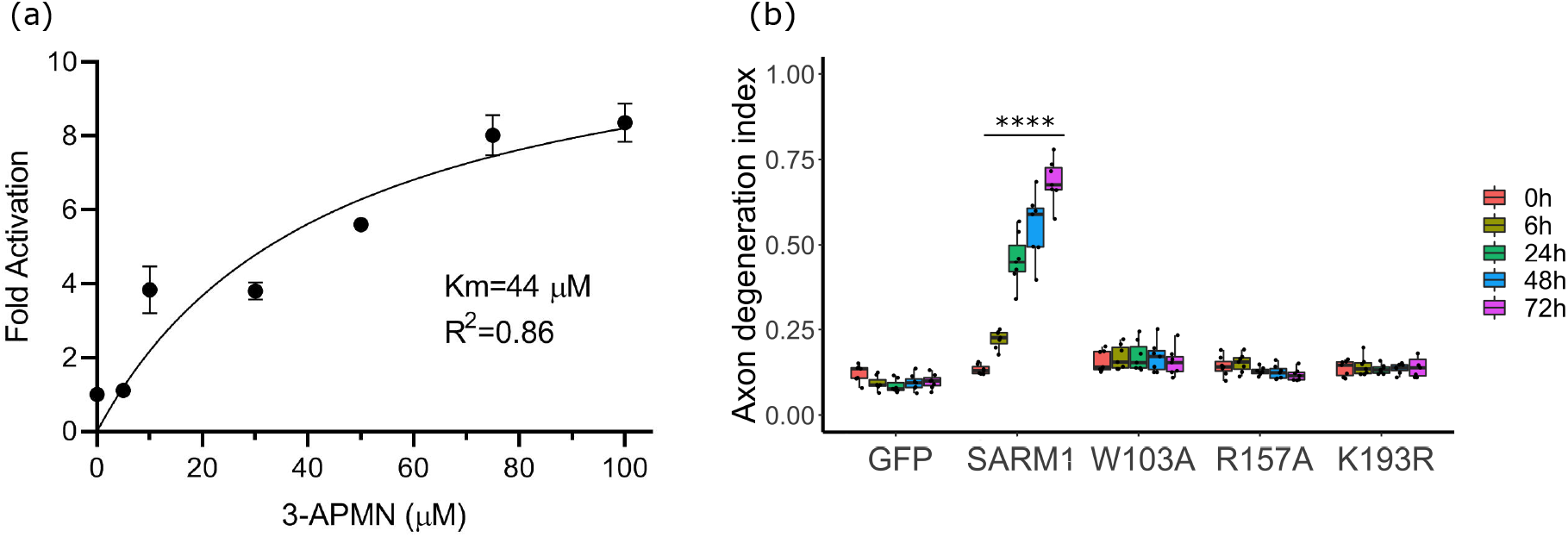
3-APMN directly activates SARM1 via its allosteric binding site. (a) Fold-increase in rate of SARM1 hydrolase activity with increasing concentrations of 3-APMN (Mean± SD, n=3), as measured by PC6 fluorescence. (b) 3-APMN activates SARM1 via its NMN/NAD^+^-binding pocket in the N-terminal domain. *Sarm1* KO DRG neurons were infected with lentivirus expressing W103A, R157A or K193R SARM1 mutants or controls (GFP and wild-type SARM1) at DIV3. Axon degeneration indexes were determined at indicated times after 3-AP (300 μM) administration. The first and third quartile, and the median are shown in boxplot with ± 1.5 time interquartile. Statistical significance was determined by one-way ANOVA tests, comparing to degeneration index of 3-AP+GFP at each time point. *p < 0.05; **p < 0.01; ***p < 0.001; ****p < 0.0001.

To determine whether 3-APMN activates SARM1 via the same binding pocket responsible for NMN-mediated SARM1 activation, we examined several SARM1 mutants in which this allosteric pocket is disrupted (Figley et al., 2021). We tested SARM1 W103A, R157A and K193R, three mutants in the allosteric pocket that are not activated in response to elevated NMN (Figley et al., 2021). Tellingly, lentiviral expression of wildtype SARM1 restores 3-AP-induced axon degeneration to *Sarm1* KO DRG neurons, whereas the axons of 3-AP-treated neurons expressing any of the three SARM1 allosteric pocket mutants remain intact (Fig. 4b). Hence, the N-terminal allosteric site that binds NMN and NAD^+^ is necessary for 3-AP-induced SARM1 activation.

### 3-AP neurotoxicity in mice requires SARM1

Early animal experiments showed that intraperitoneal (IP) injection of 3-AP leads to rapid death accompanied by extensive lesions throughout the nervous system (Desclin et al., 1974). To investigate the role of SARM1 activation in 3-AP toxicity *in vivo*, we administered 3-AP via IP injection to WT and *Sarm1* KO mice. Wild type mice treated with 275 mg/kg 3-AP generally survive, but those treated with 400 mg/kg 3-AP die rapidly. In contrast, *Sarm1* KO mice survive and have no obvious phenotype following treatment with up to 500 mg/kg 3-AP, a dose that is rapidly lethal for wild type mice (Fig. 5a). To assess SARM1 activation *in vivo*, we monitored levels of cADPR, a SARM1 biomarker. Following IP treatment with 275 mg/kg, levels of cADPR in the sciatic nerve increase 3-4 fold by 3 days post-administration (Fig. 5b). Similar injections into *Sarm1* KO mice caused no alteration in cADPR levels, confirming the SARM1-dependence of cADPR production.

**Figure 5.**
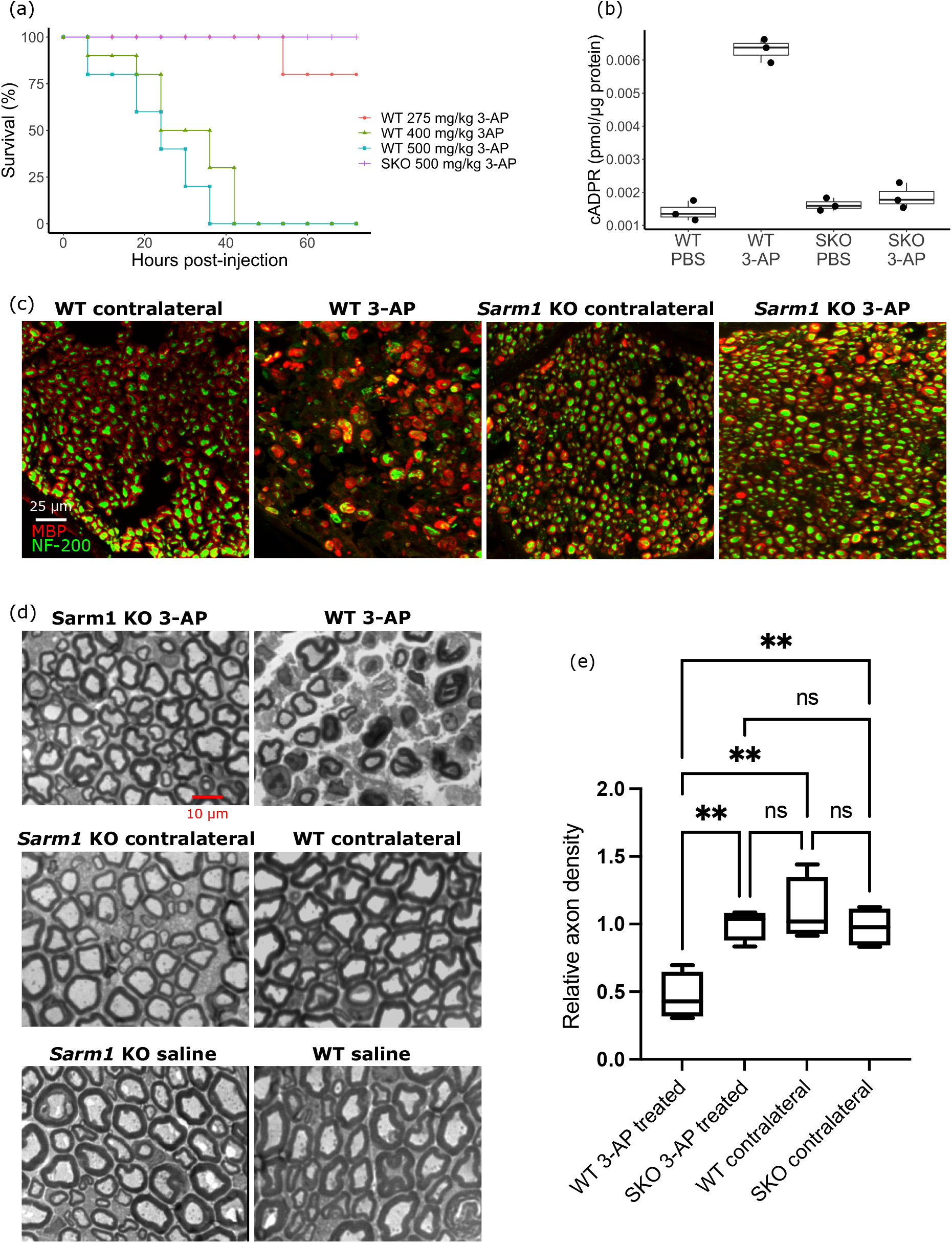
3-AP neurotoxicity in mice is mediated by SARM1. (a) Survival curve of WT and *Sarm1* KO (SKO) mice injected IP with indicated doses of 3-AP (n=5). Mice injected with PBS (control) showed no lethality (n=3), and SKO mice injected with 275 mg/kg 3-AP showed no lethality (n=5). (b) cADPR levels in sciatic nerve of WT and SKO mice at 3 days post IP injection with PBS or 3-AP (275 mg/kg). (c) Immunofluorescent staining with antibodies against neurofilament (NF-200) or myelin basic protein (MBP) of tibial nerve cross sections from WT and *Sarm1* KO animals in which the sciatic nerve was wrapped with Surgifoam soaked in 3-AP (500 mM) and corresponding contralateral untreated nerve. (d) Images of toluidine blue-stained sections of tibial nerve treated with Surgifoam soaked in saline or 3-AP (500 mM) and corresponding contralateral untreated nerves. (e) Quantification of axon counts relative to counts of saline treated nerve from toluidine blue-stained images. The first and third quartile, the median are shown in boxplot with ± 1.5 time interquartile. Statistical significance was determined by one-way ANOVA tests. *p < 0.05; **p < 0.01; ***p < 0.001; ****p < 0.0001.

To examine the effect of 3-AP on axon degeneration and to circumvent systemic toxicity, we performed local administration by applying a Surgifoam wrap soaked in 500 mM 3-AP (or saline control) to surgically exposed sciatic nerve. After such treatment, WT mice rapidly develop an abnormal gait, favoring their untreated leg, and fail to splay the toes of their 3-AP treated paw in the usual fashion. Seven days after local treatment, the tibial nerves distal to the 3-AP treatment site were dissected and analyzed. 3-AP treated nerves from WT mice showed a dramatic loss of axons compared to the contralateral nerve or saline-treated nerve, demonstrating the local effect of 3-AP. In contrast to the findings in WT mice, treated nerves from *Sarm1* KO mice were well protected from axon loss (Fig. 5c-e). Quantification of axons in the tibial nerve shows a substantial axon loss in the 3-AP treated nerves (Fig. 5e). These studies demonstrate that 3-AP triggers SARM1-dependent axon degeneration *in vivo*, and highlights that direct application of such a SARM1-activating neurotoxin is capable of triggering local axon loss.

### Additional nicotinamide analogues induce SARM1-dependent axon degeneration

The pyridine ring is a common component of many drugs and industrial chemicals, suggesting that many of these compounds could also be neurotoxic in a SARM1-dependent fashion. To test whether other pyridine derivatives also bind and activate SARM1, we tested 2-, 3-, and 4-aminopyridines and 2- 3-, and 4- acetylpyridines in our axon degeneration and SARM1 activation assays. Interestingly, we found that 2-aminopyridine causes axon degeneration in WT but not in *Sarm1* KO neurons, similar to 3-AP (Supplemental Fig. 3). Hence, multiple pyridine compounds exhibit SARM1-dependent toxicity, identifying SARM1 as a candidate target for the many toxins with related chemical structures (Buonvicino et al., 2018). Indeed, a recent preprint demonstrated that the rodenticide vacor, which has a pyridine ring, induces SARM1-dependent toxicity (Loreto et al., 2020).

## DISCUSSION

A major advance in our understanding of programmed axon degeneration was the recent discovery that SARM1, the central axon executioner, is activated by a rise in the intracellular ratio of the nicotinamide metabolites NMN to NAD^+^ (Figley et al., 2021). Here we capitalize on this breakthrough to expose the mechanism of a decades-old biological riddle, the dramatic neurotoxicity of 3-AP and related compounds (Woolley et al., 1943). Our results support a model in which 3-AP is metabolized to produce the NMN mimetic 3-APMN which binds the allosteric pocket of SARM1, thereby activating its NADase activity and initiating pathological axon degeneration. The identification of SARM1 as a direct target of neurotoxins has two important implications. First, these findings suggest that SARM1 activation by environmental toxins is a potential contributor to neurodegenerative disorders. Second, the identification of selective SARM1 agonists may lead to the development of new approaches for therapeutic neurolysis.

The nervous system employs an active program of axon self-destruction, also known as Wallerian degeneration, to facilitate the orderly clearance of axon segments damaged by trauma or disease. The choice between axon maintenance and dissolution is chiefly made by the TIR-containing NAD^+^ hydrolase SARM1 (Figley et al., 2020). Healthy neurons maintain SARM1 in an autoinhibited state, but injury- or disease-associated attrition of the NAD^+^ synthetase NMNAT2 (Gilley et al., 2010) induces rapid NAD^+^ depletion by SARM1 leading to metabolic catastrophe and axon fragmentation. Thus NAD^+^ homeostasis is crucial for axon maintenance as well as overall neuronal health (Gerdts et al., 2015). NAD^+^ is generated from Nam in two steps: the rate-limiting conversion of Nam to NMN by NAMPT, followed by synthesis of NAD^+^ from NMN by NMNAT2 and its paralogs (Sauve, 2008). NMN and NAD^+^ compete to bind an allosteric pocket that modulates SARM1 autoinhibition, rendering SARM1 a metabolic sensor that responds to an elevated NMN/NAD^+^ ratio (Figley et al., 2021; Jiang et al., 2020; Sporny et al., 2020). As such, experimental manipulations that alter the apparent concentration of either metabolite, whether by short-circuiting the usual synthesis pathway or by interjecting an NMN mimetic, directly affect SARM1 activity and its consequences (Di Stefano et al., 2017; Sasaki et al., 2009; Zhao et al., 2019).

It was first observed three quarters of a century ago that treatment with a nicotinamide analog, 3-AP, induces rapid hindlimb paralysis and death in mice, symptoms that could be prevented by sufficient prior fortification with nicotinic acid or Nam (Woolley et al., 1943). 3-AP-treated animals develop widespread nervous system lesions, particularly in brainstem nuclei, and consecutive treatment with 3-AP and Nam was eventually developed into a useful method for specific ablation of the inferior olive in rodents (Desclin et al., 1974; Rondi-Reig et al., 1997; Wecker et al., 2017). However, the mechanism of 3-AP neurotoxicity remained unknown. Here we present comprehensive evidence demonstrating that 3-AP injures the nervous system by directly activating SARM1 and inducing Wallerian degeneration.

The profound invulnerability of both *Sarm1*^-/-^ mice and *Sarm1*^-/-^ cultured neurons to 3-AP first alerted us to the SARM1-dependence of 3-AP toxicity and lead to the straightforward hypothesis that 3-APMN, a 3-AP metabolite nearly identical to NMN, hijacks the SARM1 pathway by binding the enzyme’s allosteric site and triggering its pro-degenerative activity. Analysis of 3-AP-treated wildtype neurons confirmed the presence of 3-APMN and its derivative 3-APAD, as well as the specific SARM1 activity marker, cADPR. Decisively establishing our thesis required two further key pieces of evidence: 1) the demonstration that 3-APMN directly binds and activates SARM1, and 2) that manipulating 3-APMN levels in 3-AP-exposed neurons correspondingly alters SARM1 activity and its toxic consequences. Both of these criteria were demonstrated using both *in vitro* and *in vivo* methods, elucidating the mechanism of action of this long-studied neurotoxin. Moreover, this proposed mechanism of SARM1-dependent 3-AP toxicity surely applies to other structurally-similar pyridine derivatives, and we identified 2-aminopyridine as such a case. We suggest that SARM1 activation may be a common cause of neurotoxicity caused by pyridine compounds.

In the course of demonstrating the SARM1-dependence of 3-AP toxicity *in vivo*, we exposed peripheral nerves to 3-AP in order to determine whether the toxin could stimulate localized neurodegeneration. Prior *in vivo* studies of 3-AP were confined to systemic administration. A key motivation for these experiments was to explore the potential utility of 3-AP or other pyridines for therapeutic neurolysis—the application of physical or chemical agents to effect temporary degeneration of nerves distal to a targeted lesion (Filippiadis et al., 2019). Purposeful nerve destruction can be appropriate for a variety of severe pain conditions, especially visceral pain associated with pancreatic cancer, trigeminal neuralgia, or facetogenic and vertebral pain (D’Souza et al., 2021). Gratifyingly, we found that local application of 3-AP caused rapid degeneration of the exposed nerve fibers without apparent systemic harm. Currently, chemical neurolysis is achieved by injecting phenol or ethanol, nonspecific noxious agents that can damage non-neuronal tissues. While both chemicals are highly efficacious, ethanol injection produces severe burning pain and phenol may cause systemic complications such as nausea, cardiovascular depression and cardiac arrhythmias (Weksler et al., 2007). Therefore, a neuroselective agent could expand the option of neurolysis beyond its current limited indications to more broadly treat neuropathic pain. Because neurolytic techniques specifically exploit Wallerian degeneration to ablate axons (D’Souza et al., 2021; Filippiadis et al., 2019), we propose that a SARM1 agonist fits well with this application. Thus, in the best traditions of medicine, the solution to a toxicological mystery may be adapted into a useful therapeutic.

## ACKNOWLEDGEMENTS

We thank Rachel McClarney, Simburger Kelli, Timothy Fahrner, Cassidy Menendez for technical assistance. The work was supported by National Institutes of Health (Grant RF1AG013730 (J.M.), R01CA219866 and RO1NS087632 (J.M. and A.D.)).

## AUTHOR CONTRIBUTIONS

Conceptualization, T.W., J.Z., A.S., K.W.K., Y.S., A.D., J.M.; Investigation, T.W., J.Z., A.S., K.W.K., Y.S., C.D., Y.Y., M.D.F., X.M.; Methodology, T.W., J.Z., A.S., K.W.K., Y.S.; Formal Analysis, T.W., J.Z., A.S., Y.S., C.D., Y.Y., M.D.F., X.M., A.N.; Resources, J.Z., A.S., K.W.K., Y.S., A.N.; Writing, T.W., J.B., J.Z., A.S., A.N., A.D., J.M.; Supervision, J.B., A.D., J.M.; Funding Acquisition, A.D., J.M.

## EXPERIMENTAL MODEL AND SUBJECT DETAILS

### Reagents and plasmids

The NAMPT inhibitor FK866 was obtained from the National Institute of Mental Health Chemical Synthesis and Drug Supply Program (MH number F-901). Nicotinamide (Nam), nicotinamide mononucleotide (NMN), nicotinamide adenine dinucleotide (NAD^+^), 3-acetylpyridine (3-AP), 3-acetylpyridine adenine dinucleotide (3-APAD), thionicotinamide (thio-Nam), and thionicotinamide adenine dinucleotide (thio-NAD) were obtained from Sigma-Aldrich (St. Louis, MO). For DRG neuron culture, Neurobasal Medium (Life Technologies, Carlsbad, CA) was supplemented with B27 (Invitrogen, Waltham, MA), NGF (Harlan Bioproducts, Indianapolis IN), 5-fluoro-2’-deoxyuridine (Sigma-Aldrich) and uridine (Sigma-Aldrich). 96-well and 24-well tissue culture plates (Corning) for DRG culture were coated with poly-D-Lysine (Sigma-Aldrich) and laminin (Invitrogen). Lentivirus vector constructs harboring cDNAs including mCherry, *cytNMNAT1* (*mouse*), *NAMPT* (*mouse*), *NRK1* (*mouse*), *NMN deamidase* (*E. coli*) and mutant *SARM1* were previously described (Figley et al., 2021; Sasaki et al., 2016).

### Lentivirus production

Lentivirus production was performed as described previously (Araki et al., 2004). HEK 293T cells were seeded with the density of 1 × 10^6^ in 35 mm well tissue culture plate (Corning). Transfection was performed one day after seeding the cells. Lentiviral constructs were cotransfected with VSV-G and psPAX2 lentiviral packing vectors. Two days after transfection, lentiviruses were collected from culture supernatant and concentrated with Lenti-X concentrator (Clontech). Lentiviruses were suspended in 1/10 volume of culture supernatant with PBS, and stored at −80 °C.

### DRG neuron culture

Mouse DRG culture was as described previously (Sasaki et al., 2009). Mouse DRGs were dissected at embryonic day 13.5. Approximately 50 ganglia were collected from each embryo and dissociated in 0.05% Trypsin containing 0.02% EDTA (Gibco) at 37 °C for 20 min. The DRG neurons were washed 3 times with DRG culture medium after incubation with Trypsin. Cells were seeded as spot cultures in 96-tissue culture plates (Corning) coated with 0.1 mg/ml poly-D-lysine and 0.1 mg/ml laminin. They were cultured in Neurobasal (Gibco) containing 2% B27 (Invitrogen), 100 ng/ml 2.5S NGF (Harlan Bioproducts), and 1 μM 5-fluoro-2’-deoxyuridine/1 μM uridine. The culture medium was exchanged with fresh DRG culture medium every 2-3 days. Lentiviruses were added 3 days after seeding (DIV3).

### Quantification of axon degeneration

After culture for 7 days, axotomy was performed by cutting DRG axons manually with a razor blade. After axon transection, bright-field images of distal axons were taken using the Operetta automated imaging system (PerkinElmer). Axon degeneration indexes were calculated from these bright-field images using ImageJ (NIH). The axon degeneration index was calculated as the ratio of fragmented axon areas as previously described (Sasaki et al., 2009). Indexes were calculated as the average of six fields per well.

### NAMPT and NMNAT1 production in *E. coli*

StrepTag-NAMPT and StrepTag-NMNAT1 were cloned into pET30a+ plasmids. Plasmids were transformed into T7 Express competent *E. coli* (New England BioLabs) as described previously (Essuman et al., 2017). Single colonies were selected and grown overnight at 37 °C. Cultures were diluted in LB media, IPTG (1 mM) was added at A_600_ =0.6, and bacteria were induced for another 4 hr before harvest. Protein purification was performed using StrepTactin Megbeads (Cube Biotech). The beads were washed with HEPES buffer (50 mM HEPES/NaOH pH 7.5, 300 mM sodium chloride, 20% glycerol with EDTA-free protease inhibitors) and proteins were eluted with 5 mM dethiobiotin (50 mM HEPES/NaOH pH 7.5, 300 mM sodium chloride, 20% glycerol, 5 mM dethiobiotin with EDTA-free protease inhibitors). Protein concentrations were determined by Pierce™ BCA Protein Assay (Thermofisher Scientific).

### SARM1 protein purification from mammalian cells

A HEK 293T cell line stably expressing Nicotinamide Riboside Kinase 1 (NRK1) was used to produce SARM1 protein as previously described (Essuman et al., 2017). In short, plasmid containing N-terminal StrepTag-SARM1 was transfected into NRK1-HEK 293T cells using X-tremeGENE reagent (Millipore Sigma). After 2 days, the cells were harvested and re-suspended with binding buffer (100 mM Tris-HCL pH 8, 150 mM sodium chloride, 0.01% Tween-20 with EDTA-free protease inhibitors (Pierce)). After sonication, cell debris were removed by centrifugation at 15,000 xg for 10 minutes. Soluble lysates were then mixed with StrepTactin Megbeads (Cube Biotech) equilibrated with the same binding buffer. After incubation at 4 °C for 60 minutes, beads were washed with washing buffer (25 mM HEPES, pH 7.5 and 150 mM NaCl with EDTA-free protease inhibitors). If elution was not required, SARM1 containing beads were re-suspended in washing buffer with addition of 1 mM TCEP and stored at −80 °C before further use.

### Base exchange assay and metabolite extraction for HPLC and LC-MS/MS

SARM1 protein was eluted from the beads by incubating for 15 min at 4 °C degree in eluting buffer (25 mM HEPES/NaOH pH 7.5, 150 mM sodium chloride, 20% glycerol, 5 mM desthiobiotin with EDTA-free protease inhibitors). In a typical assay, 500 ng SARM1 protein was mixed with 50 μM or 100 μM NAD^+^ and ascending concentrations of 3-AP (20 μM, 40 μM, 60 μM). The reactions were incubated at 37 °C degree for 1 hr in a total reaction volume of 50 μl. For HPLC analysis, the reaction was stopped by addition of 50 μl 0.5 M perchloric acid (HClO4) and then neutralized with 3 M potassium carbonate (K2CO3). After neuralization and removal of precipitate by centrifugation, 50 μl supernatant was loaded onto HPLC. For LC-MS/MS analysis, the reaction was stopped by addition of 50 μl 50% methanol and metabolites were extracted using 1/3 total volume of chloroform. The aqueous phase was lyophilized and stored at −20 °C for LC-MS/MS analysis.

### 3-APMN identification by LC-MS/MS and purification by FPLC

Reactions were carried out using 5 μg NAMPT enzyme, 500 μM 3-AP, 1 mM ATP, 500 μM PRPP in HEPES buffer (50 mM HEPES pH 7.5, 20 mM MgCl2) and incubated for 6 hr at 37 °C. Reactions with or without PRPP (obligate cofactor) were stopped by addition of equal volume of 50% methanol and metabolites were extracted using chloroform. The lyophilized aqueous phase was analyzed by LC-MS/MS to identify 3-APMN. 3-APAD was identified similarly from reactions containing both NAMPT and NMNAT1 using LC-MS/MS. After HPLC separation, 3-APMN was purified through FPLC (Supp. Fig. 2). In short, an ion-exchange chromatography (IEC) was carried out on AKTA Purifier (GE HealthCare) using an anion exchange column Source 15Q 4.6/100 PE (GE HealthCare). The column was equilibrated with 0.01 M ammonium acetate, pH 9.0 at room temperature with a flow rate of 1 ml/min. After injection of the reaction mixture, a linear gradient elution was applied by mixing the 0.01 M and 1 M ammonium acetate, pH 9.0 buffers at 1 ml/min. The eluate was monitored at wavelength 260 nm. Peaks were collected and lyophilized. Next, lyophilized samples were reconstituted with 5 mM ammonium formate and the identity of 3-APMN peak was confirmed using LC-MS/MS (Agilent, 6470 Triple Quad LC/MS) with a C18 column (Agilent, EclipsePlus C18, 3.0 × 50mm, 1.8mm particles).

### SARM1 activation assay by PC6 fluorescence

Modified from previous paper (Li et al., 2021), typically, in a mixture of 50 μl, ~200 ng of SARM1(on-beads) was mixed with 50 μM PC6, 50 μM NAD^+^ and 0 to 100 μM of purified 3APMN. The reaction was carried out in a Bio Tek’s Cytation 5 plate reader at 25 °C. Fluorescence signal, which corresponds to PAD6 formation, was recorded with 350 nm excitation and 450 nm emission wavelengths every 2 minutes. The production rate of PAD6 was calculated based on the fluorescence changes within the first 30 minutes of reaction.

### Metabolite measurement using LC-MS/MS

Lyophilized samples were reconstituted with 50 μl of 5 mM ammonium formate and centrifuged at 12,000 g for 10 min. Cleared supernatants were transferred to sample vials. Serial dilutions of standards for each metabolite in 5 mM ammonium formate were used for calibration. HPLC-mass spectrometry analysis was performed on an Agilent 1290 Infinity II liquid chromatography system (Agilent Technologies, Santa Clara, CA) with a flexible pump, multisampler, sample cooler and an MCT containing an Atlantis T3 column (2.1 × 150 mm, 3 μm) and VanGuard guard cartridge (2.1 mm X 5 mm, 3 μm) (Waters, Milford, MA), coupled to an Agilent 6470 Triple Quad mass spectrometer (Agilent Technologies, Santa Clara, CA). The mobile phase (0.15 ml/min) was 5 mM ammonium formate in water (A) and 100% methanol (B). The column was equilibrated with 0% B, maintained after injection for 2 min, then a linear gradient to 20% B applied over 4 min. The column was then ramped to 50% B over 2 min, and held at 50% for 2 min, then reverted back to 0% B over the next 5 min and allowed to re-equilibrate at 0% B for 9 min. The total run time was 24 min per sample. The injection volume was 10 μl. The mass spectrometer was equipped with an electrospray ion source which was operated in positive ion multiple reaction monitoring (MRM) mode for the detection of all of the metabolites. The [M+H]+ transitions were optimized for each metabolite and were selected as follows: m/z 664 → 428 for NAD^+^, m/z 335 → 123 for NMN, m/z 542 → 428 for cADPR, m/z 508 → 136 for ATP, m/z 663 → 136 for 3-APAD and m/z 334 → 122 for 3-APMN. The mass spectrometer settings for the fragmentation, the collision energy (CE) and the cell accelerator voltage were optimized for each of these transitions. Raw data were acquired and quantified using MassHunter Workstation software version B.08.00 for 6400 Series Triple Quadrupole (Agilent Technologies, Santa Clara, CA).

### IP injection and local application of 3-AP to sciatic nerve using Surgifoam wrap

5 week-old mice were used for IP injection experiment. Suitable skin incisions were made at the mid-thigh level of 5 week-old mice, followed by a small incision or blunt dissection to the muscle fascia to permit retraction of the biceps femoris and gluteus superficialis muscles to expose the sciatic nerve. The nerve was carefully separated from surrounding connective tissue and then a 0.5 X 1 cm piece of sterile absorbable hemostatic material (Surgifoam) soaked in 500 mM 3-AP (or saline control) was wrapped around the sciatic nerve. Wound clips, and/or tissue adhesive were used to close the incision.

### Tissue processing

Tibial nerves from both control and 3-AP treated sides was dissected out 7 days postsurgery. The distal 2 cm portion of the nerve was fixed in 4% paraformaldehyde for 1 hr at room temperature, then moved to 30% sucrose and embedded in OCT for cryosectioning in the following day. Immunocytochemistry was performed on tibial cryosections using NF200 (Sigma N4142, 1:1000) and MBP (Millipore MAB386, 1:100) followed by anti-rabbit-Alexa488 (1:500) and anti-rat-Cy3 (1:500). The proximal 2 cm portion of the sciatic nerves were fixed in 3% Glutaraldehyde for processing and embedding in epoxy/plastic. Toluidine blue thick sections (400-600 nm thickness) were cut and axon counting was performed using binary image analysis (Hunter et al., 2007).

## QUANTIFICATION AND STATISTICAL ANALYSIS

Sample number n represents the number of samples with independent experiments or manipulations. One-way analysis of variance (ANOVA) were used to make comparisons between samples and two-tailed t-tests were used to calculate p-value between two groups and identify statistical significance. All error bars represent standard deviation (SD). Axon degeneration indexes were calculated from distal axons in brightfield images by an ImageJ macro as described before (Sasaki et al., 2009). All line graphs and boxplots were generated by R and Graph Pad Prism 7. The top and bottom of box in boxplots represents the first and third quartiles, individually. The median is represented by a horizontal line in the box, and error bars represent 1.5 times interquartile range.

---

**Supplemental Figure 1.**
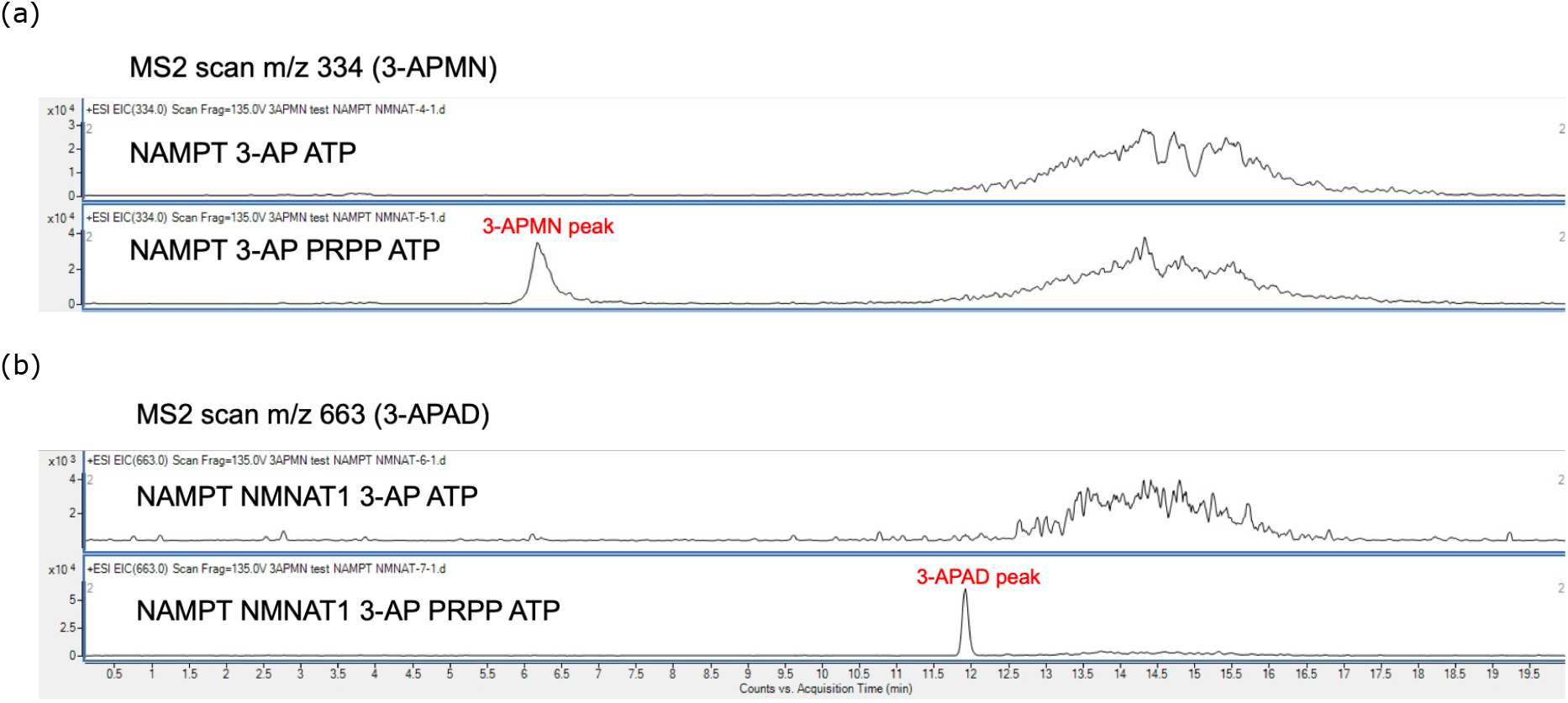
(a) 3-APMN was identified by LC-MS/MS. Purified NAMPT was incubated with ATP, 3-AP and, with or without PRPP at 37 °C for 6 hr. The reaction metabolite products were analyzed by LC-MS/MS. (b) 3-APAD was identified by LC-MS/MS. Purified NAMPT and NMNAT1 were incubated with ATP, 3-AP, with or without PRPP at 37 °C overnight. The reaction metabolite products were analyzed by LC-MS/MS.

**Supplemental Figure 2.**
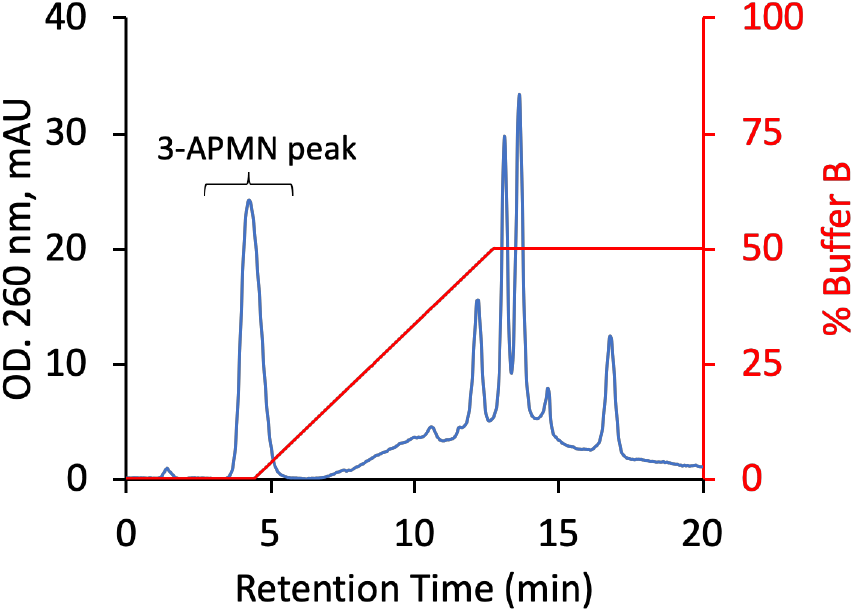
3-APMN was purified by FPLC with column Source 15Q, 4.6/100 with flow rate of 1 ml/min using Buffer B (1 M ammonium acetate, pH 9.0). The identity of the peak as 3-APMN was confirmed by LC-MS/MS.

**Supplemental Figure 3.**
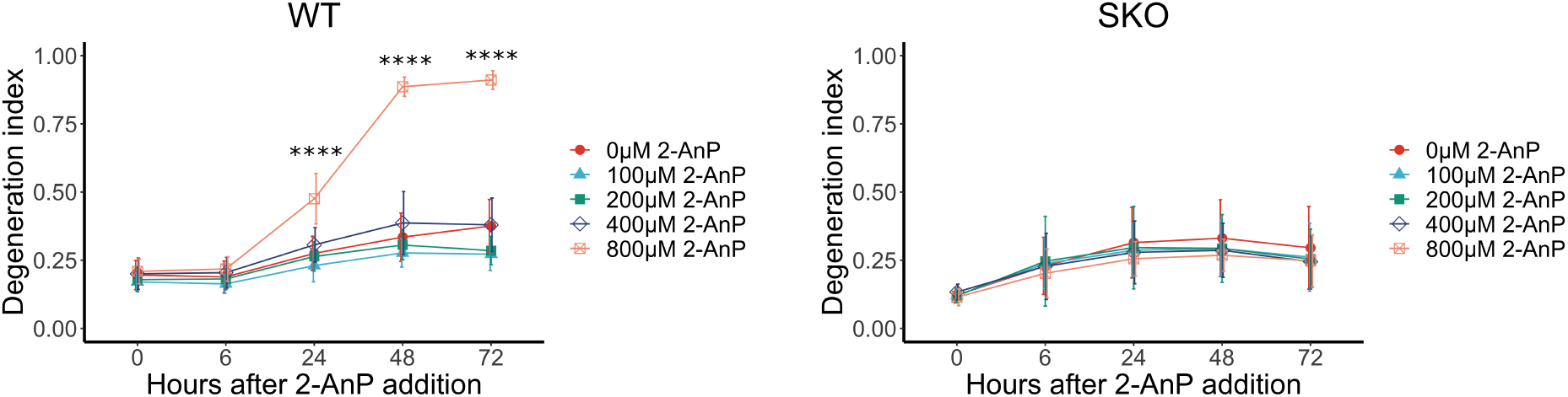
Axon degeneration index of WT and *Sarm1* KO DRG neurons treated with indicated concentrations of 2-aminopyridine (2-AnP). Statistical significance was determined by one-way ANOVA tests (compared to 0 hr). *p < 0.05; **p < 0.01; ***p < 0.001; ****p < 0.0001.

## REFERENCES

Araki, T., Sasaki, Y., & Milbrandt, J. (2004). Increased nuclear NAD biosynthesis and SIRT1 activation prevent axonal degeneration. Science. https://doi.org/10.1126/science.1098014

Beher, W. T., Holliday, W. M., & Gaebler, O. H. (1952). Studies of antimetabolites. II. 3-Acetylpyridine. The Journal of Biological Chemistry. https://doi.org/10.1016/S0021-9258(18)55512-5

Buonvicino, D., Mazzola, F., Zamporlini, F., Resta, F., Ranieri, G., Camaioni, E., … Chiarugi, A. (2018). Identification of the Nicotinamide Salvage Pathway as a New Toxification Route for Antimetabolites. Cell Chemical Biology. https://doi.org/10.1016/j.chembiol.2018.01.012

D’Souza, R. S., & Strand, N. (2021). Neuromodulation With Burst and Tonic Stimulation Decreases Opioid Consumption: A Post Hoc Analysis of the Success Using Neuromodulation With BURST (SUNBURST) Randomized Controlled Trial. Neuromodulation. https://doi.org/10.1111/ner.13273

Desclin, J. C., & Escubi, J. (1974). Effects of 3-acetylpyridine on the central nervous system of the rat, as demonstrated by silver methods. Brain Research. https://doi.org/10.1016/0006-8993(74)90627-1

Di Stefano, M., Loreto, A., Orsomando, G., Mori, V., Zamporlini, F., Hulse, R. P., … Conforti, L. (2017). NMN Deamidase Delays Wallerian Degeneration and Rescues Axonal Defects Caused by NMNAT2 Deficiency In Vivo. Current Biology. https://doi.org/10.1016/j.cub.2017.01.070

Di Stefano, M., Nascimento-Ferreira, I., Orsomando, G., Mori, V., Gilley, J., Brown, R., … Conforti, L. (2015). A rise in NAD precursor nicotinamide mononucleotide (NMN) after injury promotes axon degeneration. Cell Death and Differentiation. https://doi.org/10.1038/cdd.2014.164

DiAntonio, A. (2019). Axon degeneration: mechanistic insights lead to therapeutic opportunities for the prevention and treatment of peripheral neuropathy. Pain. https://doi.org/10.1097/j.pain.0000000000001528

Essuman, K., Summers, D. W., Sasaki, Y., Mao, X., DiAntonio, A., & Milbrandt, J. (2017). The SARM1 Toll/Interleukin-1 Receptor Domain Possesses Intrinsic NAD+ Cleavage Activity that Promotes Pathological Axonal Degeneration. Neuron. https://doi.org/10.1016/j.neuron.2017.02.022

Essuman, K., Summers, D. W., Sasaki, Y., Mao, X., Yim, A. K. Y., DiAntonio, A., & Milbrandt, J. (2018). TIR Domain Proteins Are an Ancient Family of NAD+-Consuming Enzymes. Current Biology. https://doi.org/10.1016/j.cub.2017.12.024

Figley, M. D., & DiAntonio, A. (2020). The SARM1 axon degeneration pathway: control of the NAD+ metabolome regulates axon survival in health and disease. Current Opinion in Neurobiology. https://doi.org/10.1016/j.conb.2020.02.012

Figley, M. D., Gu, W., Nanson, J. D., Shi, Y., Sasaki, Y., Cunnea, K., … Ve, T. (2021). SARM1 is a metabolic sensor activated by an increased NMN/NAD+ ratio to trigger axon degeneration. Neuron. https://doi.org/10.1016/j.neuron.2021.02.009

Filippiadis, D. K., Yevich, S., Deschamps, F., Jennings, J. W., Tutton, S., & Kelekis, A. (2019). The Role of Ablation in Cancer Pain Relief. Current Oncology Reports. https://doi.org/10.1007/s11912-019-0844-9

Gerdts, J., Brace, E. J., Sasaki, Y., DiAntonio, A., & Milbrandt, J. (2015). SARM1 activation triggers axon degeneration locally via NAD+ destruction. Science. https://doi.org/10.1126/science.1258366

Gilley, J., & Coleman, M. P. (2010). Endogenous Nmnat2 Is an Essential Survival Factor for Maintenance of Healthy Axons. PLoS Biology. https://doi.org/10.1371/journal.pbio.1000300

Hicks, S. P. (1955). Pathologic effects of antimetabolites. I. Acute lesions in the hypothalamus, peripheral ganglia, and adrenal medulla caused by 3-acetyl pyridine and prevented by nicotinamide. The American Journal of Pathology.

Horsefield, S., Burdett, H., Zhang, X., Manik, M. K., Shi, Y., Chen, J., … Kobe, B. (2019). NAD+ cleavage activity by animal and plant TIR domains in cell death pathways. Science. https://doi.org/10.1126/science.aax1911

Hunter, D. A., Moradzadeh, A., Whitlock, E. L., Brenner, M. J., Myckatyn, T. M., Wei, C. H., … Mackinnon, S. E. (2007). Binary imaging analysis for comprehensive quantitative histomorphometry of peripheral nerve. Journal of Neuroscience Methods. https://doi.org/10.1016/j.jneumeth.2007.06.018

Jiang, Y., Liu, T., Lee, C. H., Chang, Q., Yang, J., & Zhang, Z. (2020). The NAD+-mediated self-inhibition mechanism of pro-neurodegenerative SARM1. Nature. https://doi.org/10.1038/s41586-020-2862-z

Jolicoeur, F. B., Barbeau, C. M., Michele, G. de, & Barbeau, A. (1982). Influence of Nicotinamide on Neurobehavioral Effects of 3-Acetylpyridine. Canadian Journal of Neurological Sciences / Journal Canadien Des Sciences Neurologiques. https://doi.org/10.1017/S0317167100043900

Li, W. H., Huang, K., Cai, Y., Wang, Q. W., Zhu, W. J., Hou, Y. N., … Zhao, Y. J. (2021). Permeant fluorescent probes visualize the activation of SARM1 and uncover an anti-neurodegenerative drug candidate. ELife. https://doi.org/10.7554/elife.67381

Lopiano, L., & Savio, T. (1986). Inferior olive lesion induces long-term modifications of cerebellar inhibition on Deiters nuclei. Neuroscience Research. https://doi.org/10.1016/0168-0102(86)90016-7

Loreto, A., Angeletti, C., Gilley, J., Arthur-Farraj, P., Merlini, E., Amici, A., … Coleman, M. P. (2020). Potent activation of SARM1 by NMN analogue VMN underlies vacor neurotoxicity. BioRxiv.

Revollo, J. R., Grimm, A. A., & Imai, S. I. (2004). The NAD biosynthesis pathway mediated by nicotinamide phosphoribosyltransferase regulates Sir2 activity in mammalian cells. Journal of Biological Chemistry. https://doi.org/10.1074/jbc.M408388200

Rondi-Reig, L., Delhaye-Bouchaud, N., Mariani, J., & Caston, J. (1997). Role of the inferior olivary complex in motor skills and motor learning in the adult rat. Neuroscience. https://doi.org/10.1016/S0306-4522(96)00518-0

Sasaki, Y., Engber, T. M., Hughes, R. O., Figley, M. D., Wu, T., Bosanac, T., … DiAntonio, A. (2020). cADPR is a gene dosage-sensitive biomarker of SARM1 activity in healthy, compromised, and degenerating axons. Experimental Neurology. https://doi.org/10.1016/j.expneurol.2020.113252

Sasaki, Y., Nakagawa, T., Mao, X., DiAntonio, A., & Milbrandt, J. (2016). NMNAT1 inhibits axon degeneration via blockade of SARM1-mediated NAD+depletion. ELife. https://doi.org/10.7554/eLife.19749

Sasaki, Y., Vohra, B. P. S., Lund, F. E., & Milbrandt, J. (2009). Nicotinamide mononucleotide adenylyl transferase-mediated axonal protection requires enzymatic activity but not increased levels of neuronal nicotinamide adenine dinucleotide. Journal of Neuroscience. https://doi.org/10.1523/JNEUROSCI.5469-08.2009

Sauve, A. A. (2008). NAD+ and vitamin B3: From metabolism to therapies. Journal of Pharmacology and Experimental Therapeutics. https://doi.org/10.1124/jpet.107.120758

Schulz, J. B., Henshaw, D. R., Jenkins, B. G., Ferrante, R. J., Kowall, T. W., Rosen, B. R., & Beal, M. F. (1994). 3-Acetylpyridine produces age-dependent excitotoxic lesions in rat striatum. Journal of Cerebral Blood Flow and Metabolism. https://doi.org/10.1038/jcbfm.1994.134

Shen, C., Vohra, M., Zhang, P., Mao, X., Figley, M. D., Zhu, J., … Milbrandt, J. (2021). Multiple domain interfaces mediate SARM1 autoinhibition. Proceedings of the National Academy of Sciences of the United States of America. https://doi.org/10.1073/pnas.2023151118

Sporny, M., Guez-Haddad, J., Khazma, T., Yaron, A., Dessau, M., Mim, C., … Opatowsky, Y. (2020). The structural basis for SARM1 inhibition, and activation under energetic stress. BioRxiv. https://doi.org/10.1101/2020.08.05.238287

Summers, D. W., Gibson, D. A., DiAntonio, A., & Milbrandt, J. (2016). SARM1-specific motifs in the TIR domain enable NAD+ loss and regulate injury-induced SARM1 activation. Proceedings of the National Academy of Sciences of the United States of America. https://doi.org/10.1073/pnas.1601506113

Wan, L., Essuman, K., Anderson, R. G., Sasaki, Y., Monteiro, F., Chung, E. H., … Nishimura, M. T. (2019). TIR domains of plant immune receptors are NAD+-cleaving enzymes that promote cell death. Science. https://doi.org/10.1126/science.aax1771

Wang, T., Zhang, X., Bheda, P., Revollo, J. R., Imai, S. I., & Wolberger, C. (2006). Structure of Nampt/PBEF/visfatin, a mammalian NAD+ biosynthetic enzyme. Nature Structural and Molecular Biology. https://doi.org/10.1038/nsmb1114

Wecker, L., Marrero-Rosado, B., Engberg, M. E., Johns, B. E., & Philpot, R. M. (2017). 3-Acetylpyridine neurotoxicity in mice. NeuroToxicology. https://doi.org/10.1016/j.neuro.2016.11.010

Weksler, N., Klein, M., Gurevitch, B., Rozentsveig, V., Rudich, Z., Brill, S., & Lottan, M. (2007). Phenol neurolysis for severe chronic nonmalignant pain: Is the old also obsolete? Pain Medicine. https://doi.org/10.1111/j.1526-4637.2006.00228.x

Weller, M., Marini, A. M., & Paul, S. M. (1992). Niacinamide blocks 3-acetylpyridine toxicity of cerebellar granule cells in vitro. Brain Research. https://doi.org/10.1016/0006-8993(92)91043-E

Woolley, D. W., & White, A. G. C. (1943). PRODUCTION OF THIAMINE DEFICIENCY DISEASE BY THE FEEDING OF A PYRIDINE ANALOGUE OF THIAMINE. Journal of Biological Chemistry. https://doi.org/10.1016/s0021-9258(18)72240-0

Zhao, Z. Y., Xie, X. J., Li, W. H., Liu, J., Chen, Z., Zhang, B., … Zhao, Y. J. (2019). A Cell-Permeant Mimetic of NMN Activates SARM1 to Produce Cyclic ADP-Ribose and Induce Non-apoptotic Cell Death. IScience. https://doi.org/10.1016/j.isci.2019.05.001

